# Specific and non-linear effects of glaucoma on optic radiation tissue properties

**DOI:** 10.1101/2023.01.17.524459

**Authors:** John Kruper, Adam Richie-Halford, Noah C. Benson, Sendy Caffarra, Julia Owen, Yue Wu, Aaron Y. Lee, Cecilia S. Lee, Jason D. Yeatman, Ariel Rokem

**Author notes:** **Author contributions**, John Kruper, Ariel Rokem, Cecilia S. Lee, Julia Owen, Aaron Y. Lee and Jason Yeatman helped conceptualize the research ideas. John Kruper and Ariel Rokem developed and implemented the software used. Adam Richie-Halford created models for the analysis. Aaron Y. Lee, Cecilia S. Lee, and Yue Wu organized and provided the data. Aaron Y. Lee and Cecilia S. Lee provided the computing resources. John Kruper and Ariel Rokem analyzed and investigated the data. John Kruper and Ariel Rokem wrote the initial draft of the paper with input from all authors. All authors provided feedback on the manuscript. **Competing interests**, Dr. A. Lee reports grants from Santen, personal fees from Genentech, personal fees from US FDA, personal fees from Johnson and Johnson, grants from Carl Zeiss Meditec, personal fees from Topcon, personal fees from Gyroscope, non-financial support from Microsoft, grants from Regeneron, outside the submitted work; This article does not reflect the views of the US FDA.

## Abstract

Changes in sensory input with aging and disease affect brain tissue properties. To establish the link between glaucoma, the most prevalent cause of irreversible blindness, and changes in major brain connections, we characterized white matter tissue properties in diffusion MRI measurements in a large sample of subjects with glaucoma (N=905; age 49-80) and healthy controls (N=5,292; age 45-80) from the UK Biobank. Confounds due to group differences were mitigated by matching a sub-sample of controls to glaucoma subjects. A convolutional neural network (CNN) accurately classified whether a subject has glaucoma using information from the primary visual connection to cortex (the optic radiations, OR), but not from non-visual brain connections. On the other hand, regularized linear regression could not classify glaucoma, and the CNN did not generalize to classification of age-group or of age-related macular degeneration. This suggests a unique non-linear signature of glaucoma in OR tissue properties.

## Introduction

Glaucoma is the leading cause of irreversible blindness in the world ^1^. The disease causes retinal ganglion cell (RGC) death, and consequently disconnection of the transmission of visual information through the optic nerve to the lateral geniculate nucleus (LGN). The optic radiations (OR) are the brain white matter connections that further transmit the information from the LGN to the visual cortex. Though the cells in the LGN whose axons constitute the OR are not directly affected by glaucoma, they are deprived of their sensory input. A major question in sensory neuroscience, with significant clinical implications, is whether such a change to the sensory periphery affects the properties of central processing pathways ^2,3^. Examining the properties of the OR in glaucoma provides an opportunity to study the downstream effects of changes to the sensory periphery on central brain connections.

Diffusion MRI (dMRI), which measures the random motion of water within brain tissue^4^, is a non-invasive method to reconstruct the trajectory of white matter pathways, such as the OR, and to assess the physical properties of the tissue within them. DMRI has previously been used to measure the properties of white matter tissue in subjects with glaucoma ^5 6 7 8 9 10^. In several of these studies, changes were not specific to the visual pathways ^11 12 13 14^, suggesting wide-spread reorganization or systemic changes related to glaucoma that also manifest in white matter changes. One concern is that in cases where relatively small samples are used these effects could be due to uncontrolled confounds that are difficult to account for in such samples. For example, white matter tissue properties change during normal aging ^15^, and this could create a bias, because glaucoma is also more prevalent with age.

To counter subtle biases in small datasets, we can use larger datasets that have recently become more widely available. The UK Biobank (UKBB) is the largest dMRI dataset to date with many thousands of subjects^16^. In such a large dataset, confounding effects can be countered using sheer numbers, but also using statistical matching ^17^. That is, a subtle bias in the characteristics of subjects with glaucoma can be countered by creating a sub-sample of healthy subjects with closely matched age, sex, ethnicity, and socio-economic background for bias-mitigated comparison.

A powerful paradigm for detecting differences between groups – e.g., subjects with glaucoma and a statistically-matched control group – is to create machine learning (ML) systems that accurately classify these groups in a held-out dataset. ML algorithms, and particularly convolutional neural networks (CNNs)^18^, can capitalize on high-dimensional and/or non-linear patterns in data, such as the complex configuration of tissue properties along the length of a brain tract, to discriminate between different categories of individuals. They can be very accurate, overcoming significant variation across individuals, but they require large amounts of data and have also come under some criticism in their use in biomedical application, due to the “black box” nature of their operations, which can sometimes make them inscrutable ^19^. In recent years, a variety of approaches have been developed to interpret the operations of CNNs and other ML systems. For example, SHapley Additive exPlanations (SHAP) score the features that led to a decision (e.g., glaucoma or no glaucoma) in a particular individual ^20^, providing information about the decision-making process used by the ML system in a specific case, or group of cases. This approach provides both scientific interpretability, as well as potential targets for intervention.

In the present work, we used dMRI data from the UKBB to compare participants with glaucoma to a statistically matched sample of participants that do not have glaucoma. To quantify tissue properties, we used Automated Fiber Quantification^21^, an automated method, implemented in open-source software that we have developed ^22^, that delineates the OR in each individual. Tissue properties of the OR were quantified using the diffusional kurtosis imaging model (DKI)^23 24^. Statistics derived from this model are sensitive to biological changes, such as aging and disease, and, when they are used in concert, can help constrain the interpretations of the underlying biological processes ^25 26 27^. We assessed whether glaucoma effects are specific to the OR, rather than a generalized effect, by fitting ML models (CNNs) to different sub-portions of the OR that transmit information about different parts of the visual field ^28,29^, as well as to non-visual control bundles. We found that glaucoma effects are specific to portions of the OR.

One of the hypotheses about the effects of glaucoma on the white matter is that it represents accelerated aging, at least within the retina^30^. Therefore, One of the hypotheses about the effects of glaucoma on the white matter is that it represents accelerated aging, at least within the retina^30^. Therefore, we compared glaucoma effects to the effects of aging, and the effects of other retinal diseases, by comparing the generalization performance of a CNN trained to classify glaucoma in classifying: (1) subjects from different age groups; and (2) subjects with age-related macular degeneration (AMD). We also tested the converse: whether an age-discriminating CNN would accurately classify subjects with glaucoma. We found that neither CNN generalized across datasets. We used the average SHAP values to describe the differences in the features between the glaucoma-classifying CNN and the age-group classifying CNN, to further demonstrate that the pattern of changes in the OR due to glaucoma is not simply a pattern of accelerated aging.

## Results

### Statistical Matching

Statistical matching was used to create a dataset where bias due to age and other factors is negligible (Figure 1). This dataset was derived from an initial dataset with 905 subjects with glaucoma and 5,292 control subjects. In the full sample, glaucoma subjects tend to be older (*μ* ± *σ* for glaucoma: 68 ± 7; control: 62 ± 7). After Mahalanobis distance matching ^31^, we created the matched dataset A with 856 glaucoma subjects and 856 control subjects of similar ages (Figure 1b; glaucoma: 68 ± 6, control: 68 ± 6). While Figure 1 only shows age matching, we simultaneously matched on sex, ethnicity, and TDI, with similar results (Supplementary Figure 1).

**Figure 1:**
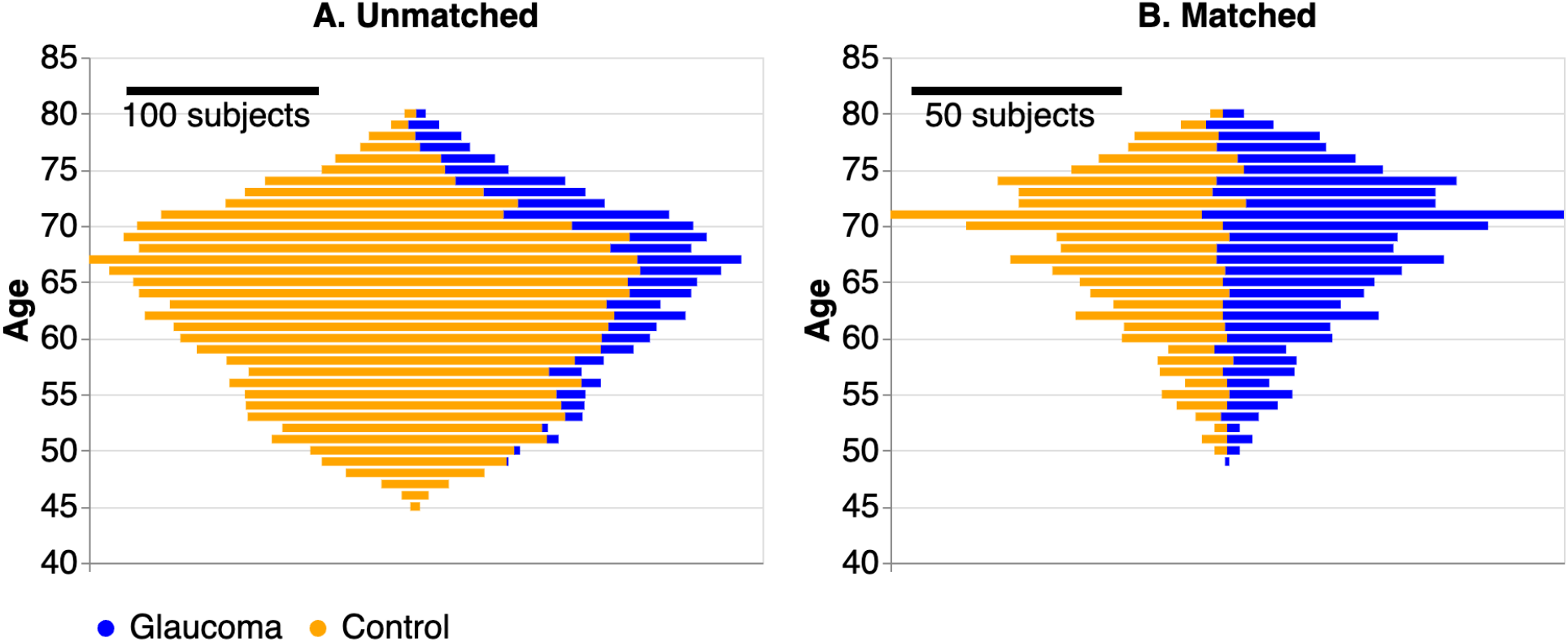
Age distribution of subjects, colored by glaucoma status. The left panel shows the entire dataset before matching. There are significantly more controls (in orange) than subjects with glaucoma (scale bars are left: 100 subjects, right: 50 subjects). Additionally, the glaucoma subjects tend to be older. The right panel shows the dataset after matching each subject with glaucoma to a control subject with the same or similar age, sex, ethnicity, and TDI.

Matching was also performed for other datasets. For datasets A.1 (age-related macular degeneration (AMD) participants, and their matched controls, see Methods) and A.2 (70-year olds, and matched 60-year olds, see Methods), 78 and 166 pairs were found, respectively. For dataset B (70-79 year olds, with matched controls 10 years younger, see Methods), 819 pairs were found. For datasets B.1 (glaucoma participants 60-64 years old, see Methods) and B.2 (AMD, see Methods), 142 and 80 pairs were found, respectively.

In dataset A, the glaucoma subjects have lower FA and MK and higher MD than the control subjects (Figure 2). However, while the mean tract profiles differ, the distributions are highly overlapping. Additionally, there are different mean tissue properties in the UNC and CST. Differences in mean tract profiles are small and not localized to the visual system.

**Figure 2:**
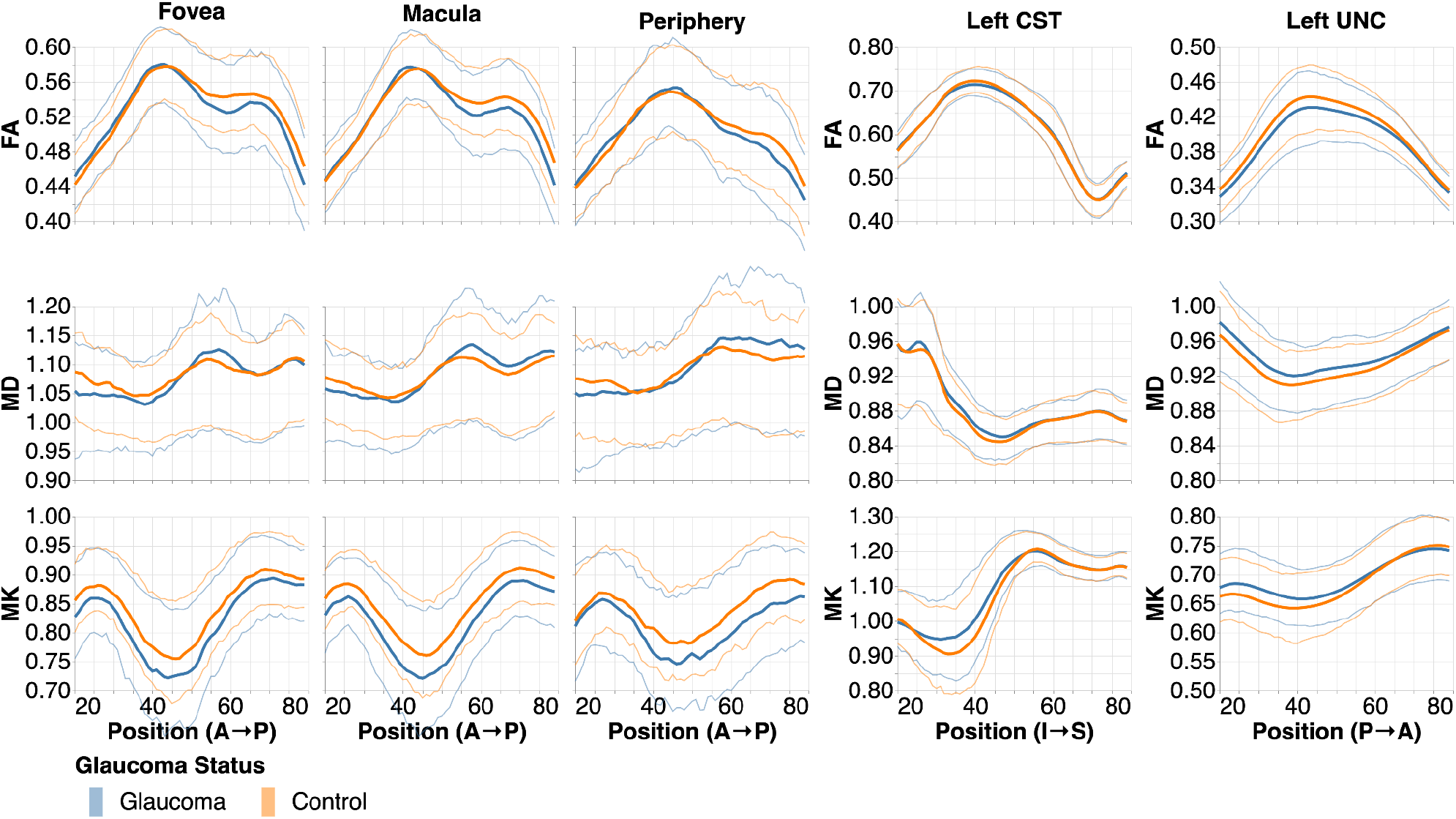
Mean tract profiles in the left hemisphere of all bundle and tissue property combinations, with thin lines showing interquartile ranges. Positions in OR bundles are from anterior to posterior (A→P), in CST are from inferior to superior (I→S), and in UNC from posterior to anterior (P→A).

### Machine Learning

Despite the high overlap between the distributions, CNNs trained on the tissue properties of white matter bundles in dataset A can accurately classify participants, but only using the tissue properties from some of the bundles (Figure 3). The ROC curves clearly separate into two groups: (1) fOR and mOR bundles, which more accurately predict glaucoma than (2) pOR and the two control bundles. We find that fOR has a significantly higher AUC than CST (DeLong’s test: p=0.0317) and UNC (p=0.0339), and mOR has significantly higher AUC than pOR (p=0.0317), CST (p=0.0119), and UNC (p=0.0119). This indicates that the tissue properties of the fOR and mOR bundles (but not the other bundles examined here) differ in participants with glaucoma compared to controls.

**Figure 3:**
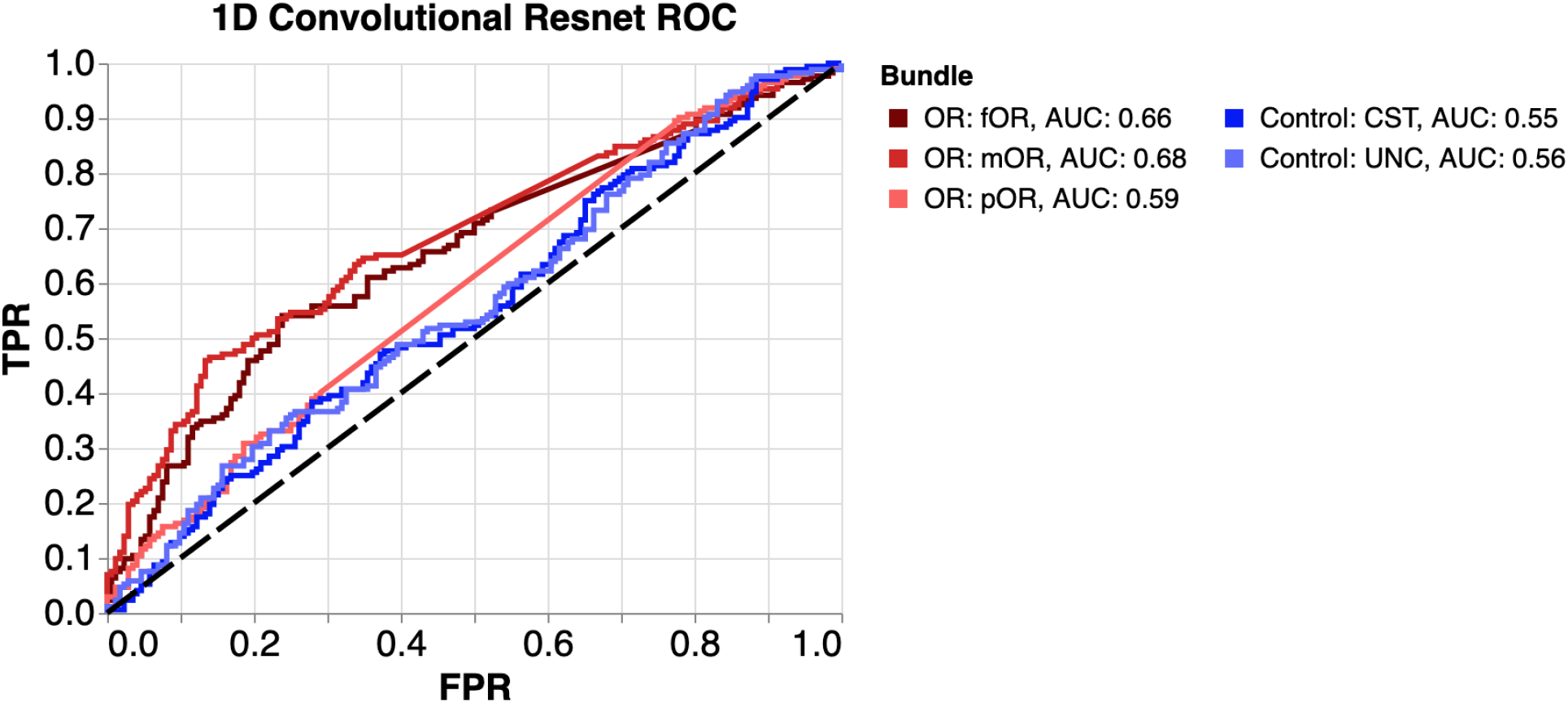
Receiver operating characteristic (ROC) curves for prediction of glaucoma using the five neural networks, each trained using tissue property from a different bundle. ROC curves compare true positive rate (TPR) to false positive rate (FPR) at various thresholds. Area under curve (AUC) is the summary of performance across all classification thresholds. The OR bundles in red tend to have better performance than the control bundles.

All bundles were not found in all subjects. We calculated, including subjects from all datasets, the extant bundle percentage for each bundle: left UNC 100.0%, right UNC 100.0%, left CST 99.8%, right CST 99.8%, left fOR 81.7%, right fOR 83.7%, left mOR 85.8%, right mOR 81.1%, left pOR 77.7%, and right pOR 67.1% (Supplementary Figure 2). We created an equivalent ROC using only test subjects where all bundles are found, which shows similar effects (Supplementary Figure 3).

As a control, we used a model of substantially less complexity than a CNN. We trained five *logistic* ridge regression models (Figure 4). If regression models can predict glaucoma with an accuracy comparable to CNNs, this would indicate that the relationship between glaucoma and the tissue properties is not complex enough to merit the use of a CNN. None of the AUC from logistic regression are significantly different from chance performance and none are significantly different from each other (Figure 4). This demonstrates that the higher AUC found from the CNN cannot be discovered with a typical linear model, further demonstrating the complexity of the relationship between glaucoma and OR tissue properties.

**Figure 4:**
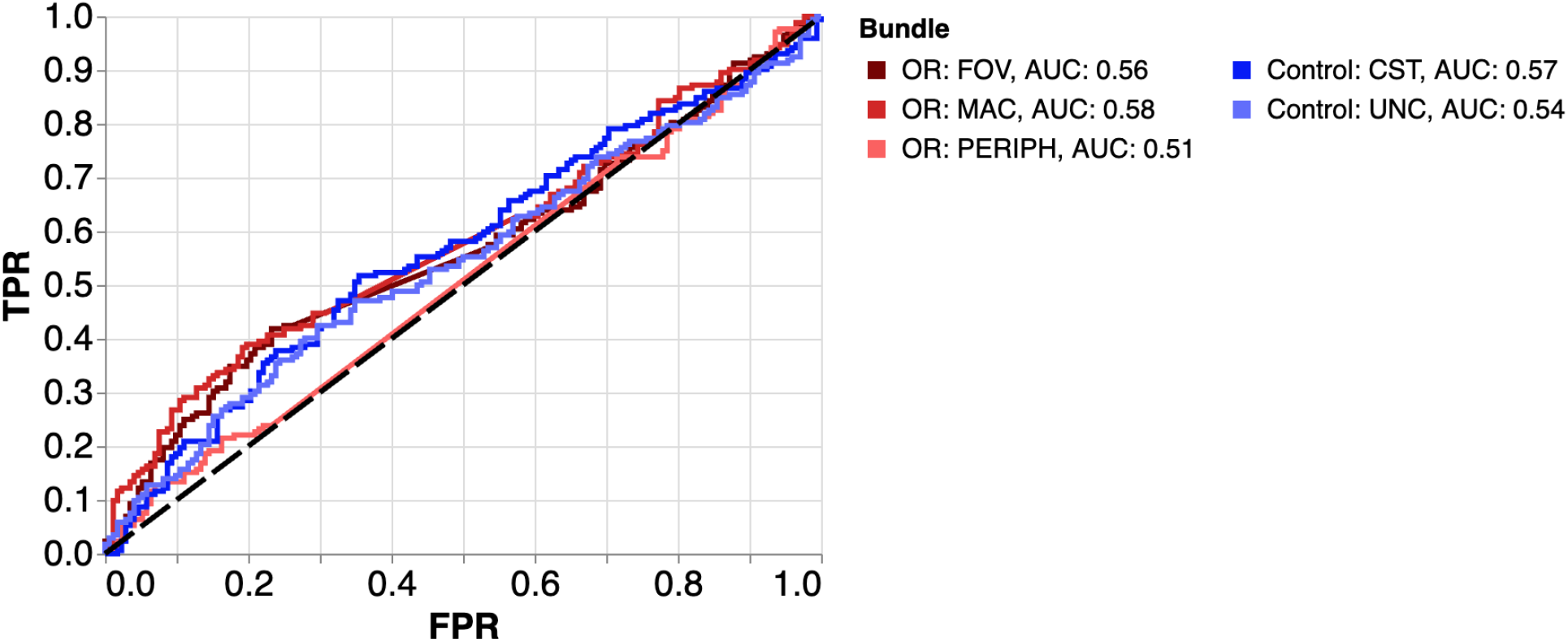
ROC curves for the prediction of glaucoma using logistic regression. Note that control bundles and OR bundles have similar AUC when using only logistic regression.

To test the generalization of our main results, we applied the glaucoma-trained network to datasets labeled according to AMD and age (datasets A.1, A.2). We found no significant AUCs (no generalization; Figure 5). Additionally, we trained five CNNs (one per bundle; the same bundles as used previously) on dataset B to classify age-group (70 vs. 60 year old). The AUCs for this network in the age-classification task are all between 0.58 and 0.63 (fOR: 0.58, mOR: 0.6, pOR: 0.6, CST: 0.63, UNC: 0.6). However, none of the bundles have AUC significantly different from each other (p>0.05 for all comparisons using the DeLong’s test), demonstrating that, unlike glaucoma, age correlates with tissue properties of the UNC and CST. Finally, we used the age-classification network to predict glaucoma and AMD (datasets B.1, B.2). Here we found no significant AUCs in the glaucoma set (AUC, fOR: 0.52; mOR: 0.58; pOR: 0.52; CST: 0.52; UNC: 0.53), but in the AMD set, there were significant AUCs for CST (0.37) and mOR (0.38).

**Figure 5:**
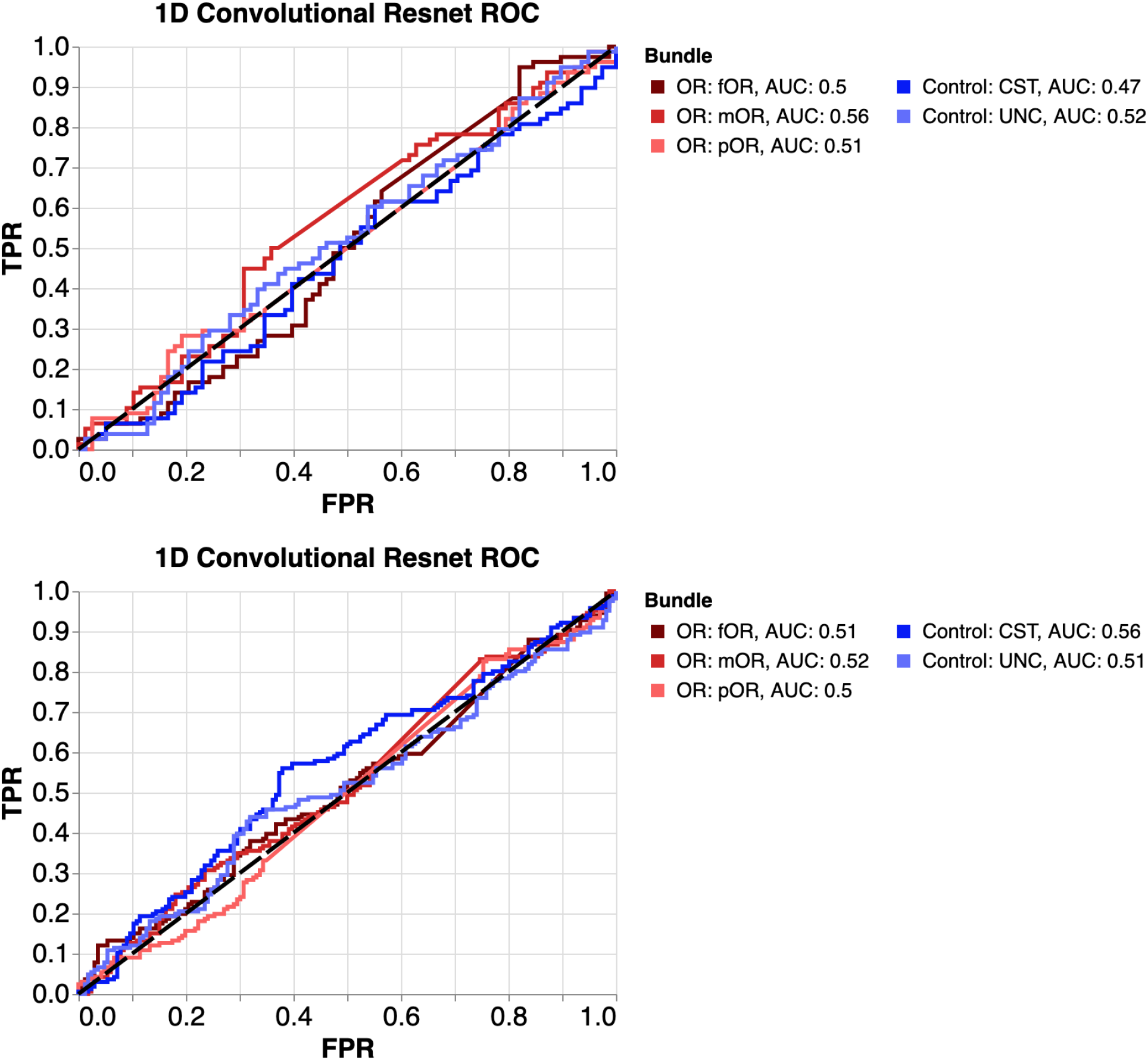
ROC curves for prediction of AMD (top) and age (bottom) using the CNN trained on the glaucoma dataset. No classifications are significantly different from chance.

We use SHAP values^20,32^ to interpret the relationship between OR tissue properties and the CNN classifications in datasets A and B using the mOR CNN (Figure 6). SHAP values use a game theoretic approach to explaining the outputs of machine learning. The 95% confidence intervals on the SHAP values from the two different datasets are often non-overlapping. This shows that the CNNs are different in terms of how they integrate information. For example, we see that higher MD indicates lower propensity to classify a participant as having glaucoma, but a higher propensity of classifying this participant as older. We also find that low FA and low MD indicate higher propensity to classify a participant as having glaucoma.

**Figure 6:**
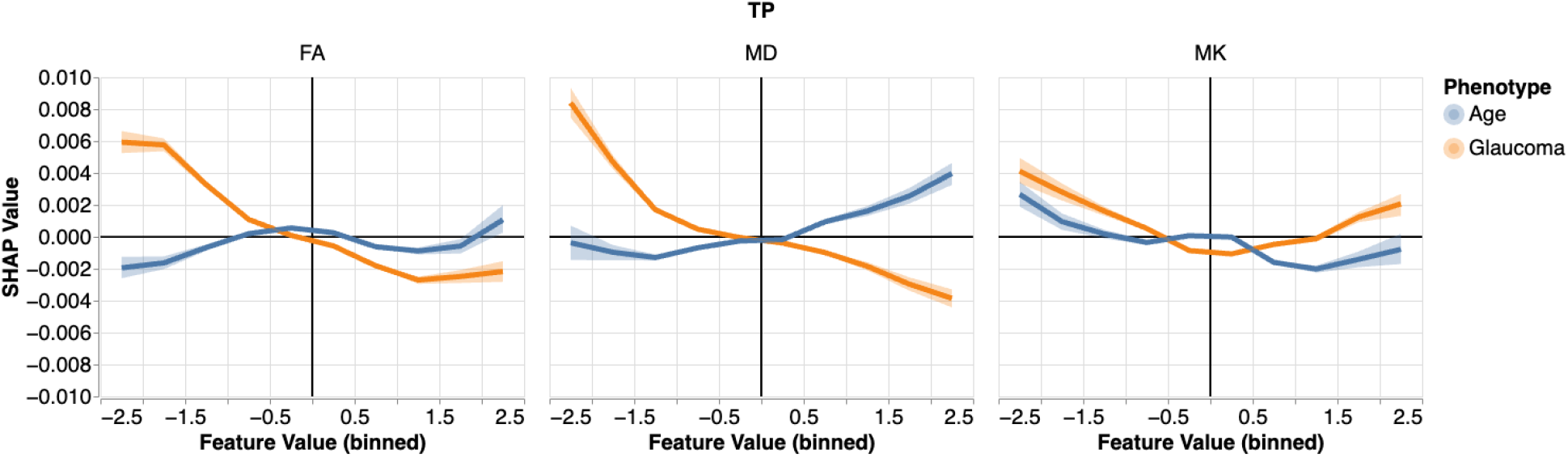
Mean SHapley Additive exPlanations (SHAP) value for each input feature value from the mOR CNN in two separate tasks. Colors indicate the dataset used, where blue is the age-group classification network on age-classification subjects (dataset B), and orange is the glaucoma-group classification network on glaucoma subjects (dataset A). Higher SHAP values indicate that a feature contributes towards predicting that the subject either has glaucoma or is older (depending on the task, blue for age group classification, orange for glaucoma classification). Columns indicate different tissue properties (FA, MD, MK), and the shading represents the 95% CI across subjects and nodes within the mOR. The x-axis shows the z-scores of each tissue property, because that is how they are provided to the neural network.

## Discussion

To study the effects of glaucoma on tissue properties in the white matter of the optic radiations, we used a large dataset of diffusion MRI measurements of participants with glaucoma from the UK Biobank ^16^, together with combinatorial methods to generate tightly-matched samples, automated methods for delineation of white matter tracts, and machine learning techniques. Previous research on glaucoma has included samples with a large diversity of characteristics ^5 6 7 8 9 10^. We sub-sampled from the UK Biobank dataset to create a group of controls that closely matched 856 participants with glaucoma in age, sex, and socioeconomic status, focusing only on these 856 pairs of participants for our analysis. This approach allowed us to mitigate potential sampling biases that could confound the conclusions about how common the effects of glaucoma in the white matter are. We found small differences in tissue properties in the optic radiations, as well as in non-visual control tracts, with large overlaps in the distributions between participants with glaucoma and the tightly-matched control group. However, using convolutional neural networks, we found that accurate classification of glaucoma participants is possible using parts of the optic radiation that represent the central visual field but not the non-visual control bundles or the parts of the optic radiations that represent the peripheral visual field. Furthermore, a linear model—regularized logistic regression—was not able to accurately classify glaucoma.

To test whether the effects of glaucoma on the white matter of the OR represent accelerated aging^30^, we cross-tested the glaucoma classification CNN in classification of subjects that differ by 10 years of age (but are matched on sex and TDI). We chose the ages 60 and 70, based on our previous observations that this is an age difference at which OR tissue properties change substantially^33^, and also the age at which glaucoma risk increases substantially in the UKBB sample ^34^. Nevertheless, the CNN trained to accurately classify glaucoma did no better than chance in classifying subjects from different age groups. In addition, a CNN trained to accurately perform this age classification task could not accurately distinguish participants with glaucoma from their matched controls. This, in addition to a separate SHAP analysis of the age-group classification CNN, suggests that the features of the OR affected in glaucoma are different from those that are affected in the normal process of aging.

To better understand these findings, we sought to interpret the performance of the CNNs. This is because, although CNNs provide accurate classification, they are difficult to interpret. We used SHAP analysis^20^, in order to describe the features that contribute to the accurate classifications by the CNNs. We found that the same features can have quite different SHAP values in the glaucoma and age classification CNNs. This, in addition to the failure of the CNNs to generalize between glaucoma and age, suggests that the features of the OR that are correlated with affected in glaucoma are different from those that are correlated with the normal process of aging.

Differences between the OR of participants with glaucoma and healthy controls could instead arise for many other reasons. One hypothesis is that the altered visual input due to the disease causes reorganization of the tissue responsible for downstream processing steps ^35–37^. These downstream changes could also be related to transsynaptic degeneration through the LGN that can be measured through changes to the OR tissue properties. Indeed, previous research found that participants with glaucoma have significantly reduced LGN volumes ^38^.

One of the puzzling findings in the present study is that accurate classification of glaucoma relies on data from the parts of the OR that represent the central visual field, and not the peripheral visual field. This is in contrast to the effects of glaucoma in the retina, which usually progress from the periphery to the center, initially causing so-called “tunnel vision.” Although this difference is less clear in subjects with no missing bundles, shown in Supplementary Figure 3, meriting caution in over-interpreting this finding. Nevertheless, functional MRI (fMRI) studies of glaucoma also found changes to the amplitude of the fMRI signal in central visual field ^36^, as well as increased cortical magnification in the central, but not peripheral, visual field in the early visual cortex of participants with glaucoma ^35^, suggesting that these changes reflect a process of remapping of functions from the peripheral visual field to the center. This reorganization at the level of the visual cortex could be a consequence of reorganization at earlier stages of the system. Taken together, our results may reflect these processes of compensation, evident in visual responses, rather than a process of degeneration.

Importantly, the present study is purely correlational, and though we have accounted for some of the obvious confounding factors (such as age), we may have not accounted for all of the differences between participants with glaucoma and controls. For example, in previous work, we found that cardiovascular fitness variables are important features in classifying participants with glaucoma in the UK Biobank dataset ^34^. In this dataset, we tried matching on cardiovascular variables but found that they did not change our results (not shown). Still, the effects measured in the OR and delineated with the CNN analysis could still reflect common underlying causes, rather than direct effects of glaucoma on the tissue properties of the OR.

Another limitation of the present study is that glaucoma and AMD status were determined based on self-report. This raises concerns that some of the individuals identified as having glaucoma have another eye disease, or no eye disease (false positives) and that there may be individuals with glaucoma among the controls (false negatives). One way in which false positives could affect our results is by the inclusion of people with other sources of sight impairments in the glaucoma group, and some of these could be impairments with a source directly in the OR. However, in our previous work with the UK Biobank dataset, we verified that self-report is highly congruent with ICD-10 diagnostic codes of glaucoma and that there was high repeatability of self-report among participants who reported glaucoma ^34^. Furthermore, a previous study that examined glaucoma in the UK Biobank dataset ^39^ verified that the distributional properties of self-reported glaucoma match the distributional properties of glaucoma patients in several other population studies ^40–42^.

Finally, there are limitations associated with the MRI methodology used. Due to the tract’s high curvature, narrow sections, and intersections with other tracts, some OR bundles were not detected in some subjects. Additionally, dMRI is unable to differentiate feedforward and feedback projections within the OR, so our conclusions encompass both. Nevertheless, the advantage of automated tractometry is clear: our large sample size allowed us to train neural networks and accurately classify glaucoma.

In summary, we used neural networks to determine that there is a complex relationship between the tissue properties of the parts of the optic radiations representing the central 14 degrees of eccentricity of the visual field. These differences persist even when participants with glaucoma are closely matched to controls. We did not find this relationship in non-visual control bundles or the parts of the optic radiation representing the peripheral visual field. This relationship is not reflective of an accelerated aging process, but may instead reflect a change in the visual input and subsequent reorganization of the visual system.

## Methods

### Datasets and Statistical Matching

We used MRI data that is available from the UK Biobank ^16^. From the available data, we created several distinct datasets, as described below. To be included in any of these, study participants must have had a dMRI data acquisition and a final visual acuity logMAR (log of the Minimum Angle of Resolution) of less than or equal to 0.3 if measured (from UKBB data field 5201).

Dataset A was composed of the following sub-samples of the UKBB data:

1. Glaucoma sub-sample: we first selected 905 subjects classified as having glaucoma (in at least one eye) by the UK Biobank’s Assessment Centre Environment (ACE) touchscreen question: “Which eye(s) are affected by glaucoma” (see UKBB data field 6119).
2. Control sub-sample: We selected 5,292 UKBB subjects for a control pool ^16,43 44^. These subjects answered “no eye problems/disorders” to the ACE touchscreen question: “Has a doctor told you that you have any of the following problems with your eyes? (You can select more than one answer)” (see UKBB data field 6148).

To reduce bias from age and other potential confounders, we use statistical matching ^17^ to create a “matched dataset”. We calculated the Mahalanobis distance ^31^ between the confounders of all pairs of glaucoma and control subjects. The confounders^45^ we used were age, sex, ethnicity, and the Townsend deprivation index (TDI)^46^. We used the ‘linear_sum_assignment’ method from Scipy 1.8.0 ^47^ to match test and control subjects such that the total Mahalanobis distance between them is minimized. This is a modified implementation of the Jonker-Volgenant algorithm ^48^. We then thresholded the matched dataset, only keeping matched subjects with a Mahalanobis distance of 0.3 or less.

After matching was concluded, we divided the resulting dataset – ***dataset A*** – into two groups: train (80%) and test (20%). From the train set, 20% was set aside as a validation set for hyperparameter selection. Matched control and glaucoma subjects were assigned into these groups in tandem. All decisions about model architecture and hyperparameters were made using dataset A training and validation sets. The results shown in this paper are from test sets, which were not viewed until final determinations about model architecture and hyperparameters were made.

To additionally test generalization of the glaucoma model, we constructed two additional *test datasets*. These datasets used control subjects that were not included in the glaucoma-matched pool (N=4,456):

1. AMD dataset (***dataset A.1***): Subjects were marked as having AMD based on the ACE touchscreen question in UKBB data (field 6148) described above. A set of matched controls was selected using statistical matching on the same criteria as above. This dataset had 81 participants with AMD and 81 matched controls.
2. Aging dataset (***dataset A.2***): 70 year-old subjects were selected from the control pool and were matched to other subjects in the control pool, with the matching modified to match to subjects 10 years younger, in addition to the other criteria used above. This dataset had 166 exactly 70-year old subjects and 166 approximately 60-year old subjects.

Another dataset – ***dataset B*** – was sub-sampled from the UKBB to train age-group classification models. Here, we selected subjects with ages 70-79 (N=962) from the control sub-sample. These were matched to other controls with the matching modified to match to subjects 10 years younger, in addition to the other criteria used above. This dataset was also divided into train, test, and cross-validation the same way as dataset A.

To test the generalization from age-group classification models, we constructed two additional *test datasets*. Controls were taken from a control pool where subjects selected for the dataset used to train the age-group classification models were removed (N=4,473):

1. Glaucoma (***dataset B.1**;* N=147). In this dataset, only participants aged between 60-64 were included.
2. AMD (***dataset B.2**;* N=81). Given lower prevalence of AMD in the overall dataset, we did not use an age inclusion criterion.

Though there is some overlap between the training data used in dataset A and in dataset B, there is no overlap between the training data and the test data in either of these datasets.

### MRI acquisition

We used preprocessed dMRI provided through the UKBB. The acquisition protocol is already described elsewhere ^44^. Briefly, dMRI measurements were conducted with a spatial resolution of 2-by-2-by-2 mm^3^, TE/TR = 14.92/3600 msec. With anterior-to-posterior phase encoding, there are five volumes with no diffusion weighting (b=0), and 50 volumes each with b=1,000 s/mm^2^ and b=2,000 s/mm^2^. An additional 6 b=0 volumes were acquired with posterior-to-anterior phase encoding direction for EPI distortion correction. Preprocessing is also described elsewhere ^43^. Briefly, the FSL “eddy” software was used to correct for head motion and eddy currents, including correction of outlier slices. Gradient distortion correction was performed and non-linear registration using FNIRT was calculated. The FNIRT mapping was also used to map the individual subjects data to the MNI template.

### Automated fiber quantification

AFQ ^21^ is an analysis pipeline that automatically delineates major white matter pathways and quantifies the properties of white matter tissue along the length of these major pathways. We used a Python-based open-source implementation of this pipeline (pyAFQ; https://github.com/yeatmanlab/pyAFQ)^22^ to extract the tissue properties of OR sub-bundles and two control bundles – the corticospinal tract (CST) and the uncinate fasciculus (UNC) – that are not a part of the visual system. In AFQ, white matter pathways are identified from a candidate set of streamlines based on anatomical landmarks: inclusion, exclusion, and endpoint regions of interest (ROIs) that are based on the known trajectory of the pathway (e.g., OR). These ROIs were transformed into each subject’s diffusion MRI coordinate frame using the FNIRT non-linear warp provided by UKBB. For UNC and CST, we also used a population-based probabilistic atlas ^49^. We seeded a GPU-accelerated residual bootstrap tractography algorithm ^50^ with 64 seeds uniformly distributed in the voxels of the inclusion ROIs to generate candidate streamlines which may follow the trajectory of the OR in each subject’s data. These candidate streamlines were filtered to recognize the OR with these conditions: streamlines (1) do not pass through the sagittal midline of the brain; (2) have at least one point that is within 3 mm of both of the inclusion ROIs; (3) do not have any point that is within 3 mm from the exclusion ROI; (4) terminate within 3 mm of the two endpoint ROIs (one in the thalamus and the other in V1)^51^. They were further filtered using the standard AFQ cleaning to remove outliers in length and trajectory ^21^. We further divided the OR into three sub-bundles based on their termination in ROIs in V1 generated from the retinotopic prior of Benson and Winawer ^52^ masked with the V1 location in the AICHA atlas ^53^. The OR sub-bundles are the foveal (fOR; terminating at parts of V1 responding to <3° degrees visual angle), macular (mOR;>=3°,<7° degrees visual angle), and peripheral (pOR; >7° degrees visual angle). Control bundles were generated using the same pipeline as the OR, using inclusion ROIs ^21^ and excluding streamlines that cross the midline.

One-dimensional tract profiles were computed for each tissue property along each of these bundles. The mean diffusivity (MD), fractional anisotropy (FA) and mean kurtosis (MK) tissue properties were calculated using the diffusional kurtosis model implemented in DIPY (https://github.com/dipy/dipy) ^54 24^. These tract profiles are computed at 100 points along each bundle, however we excluded the first and last 10 nodes, because their measured properties are more strongly influenced by partial volume effects with the gray matter. Further visualizations and analyses with machine learning use these one dimensional tract profiles.

### Machine Learning

We trained 1D convolutional residual neural networks ^55 56^ on the tract profiles from the three optic radiation bundles (fOR, mOR, pOR) and two control bundles (CST, UNC) to predict whether a patient has glaucoma. We trained one network per bundle for a total of five networks. The middle 80 nodes out of the 100 from the tract profiles were used. Thus the input to these networks consisted of 1D tract profiles of length 80 and with six channels, one for each tissue property (FA, MD, and MK) and each hemisphere (left and right). Tissue properties were normalized by converting to Z-scores before use in the neural network. In subjects where some bundles were not found, missing bundles were imputed using the mean profile of that bundle for each tissue property, separately in the train and test data^57^. We used the game-theoretical Shapley additive explanation (SHAP) values ^32^ to interpret the results of the neural network. SHAP values were calculated for each node and normalized tissue property using the SHAP software (https://shap.readthedocs.io/). To visualize SHAP values, we first divided the OR into four sections, which were the anterior and posterior segments of each hemisphere. We averaged the normalized tissue property value for all nodes in each of those four sections.

Independently, we trained 5 L2-regularized logistic regression models on the bundles. We used the liblinear solver in Scikit Learn 1.0.0 to fit these models ^58,59^. To reduce overfitting seen in the validation set, we only used every other node from the 20th to 78th node out of 100 nodes before testing. We also used the validation set to determine the level of regularization.

We applied the networks trained on dataset A to datasets A.1 and A.2. Finally, we trained five more neural networks of the same architecture on the same bundles, using dataset B. We also tested its generalizability on its corresponding B.1 and B.2 datasets.

The results of the CNNs and logistic regression are compared using areas under curve (AUC) from a receiver operating characteristic (ROC) curve ^60^. Depending on the curve, we ask one of two questions: (1) For the visual system tests (datasets A and B), are the OR bundle AUCs significantly different from the control AUCs and from each other, and (2) for the generalization tests (datasets A.1, A.2, B.1, B.2), are any of the AUCs significantly above chance. For question 1, we compare the AUC of each OR bundle to each control bundle, and the AUCs of the OR bundles to each other, resulting in nine comparisons. We calculate a p-value from the DeLong test ^60^ and then correct for the nine comparisons using the False Discovery Rate (FDR) p-value correction ^61^. We call the AUCs significantly different if the corrected p-value is below 0.05. For question 2, we use the variance calculated by the DeLong formula ^60^ to make a 95% confidence interval. We apply the Bonferroni correction to the confidence intervals ^62^ to correct for our testing of five different bundles. If the corrected confidence intervals do not include an AUC of 0.5, we call the AUC significant. The more conservative Bonferroni correction is used to correct for multiple comparisons in this case, because it is applied to confidence intervals, rather than to p-values.

### Software

All code to reproduce the analysis and the figures is available at <link to be added upon article acceptance>.

## Data Availability Statement

This study uses publicly available data from the UK Biobank. More information on the data and access can be found here: https://www.ukbiobank.ac.uk/enable-your-research

