## Supplementary Figure for "Specific and non-linear effects of glaucoma on optic radiation tissue properties"

### Supplementary Figures

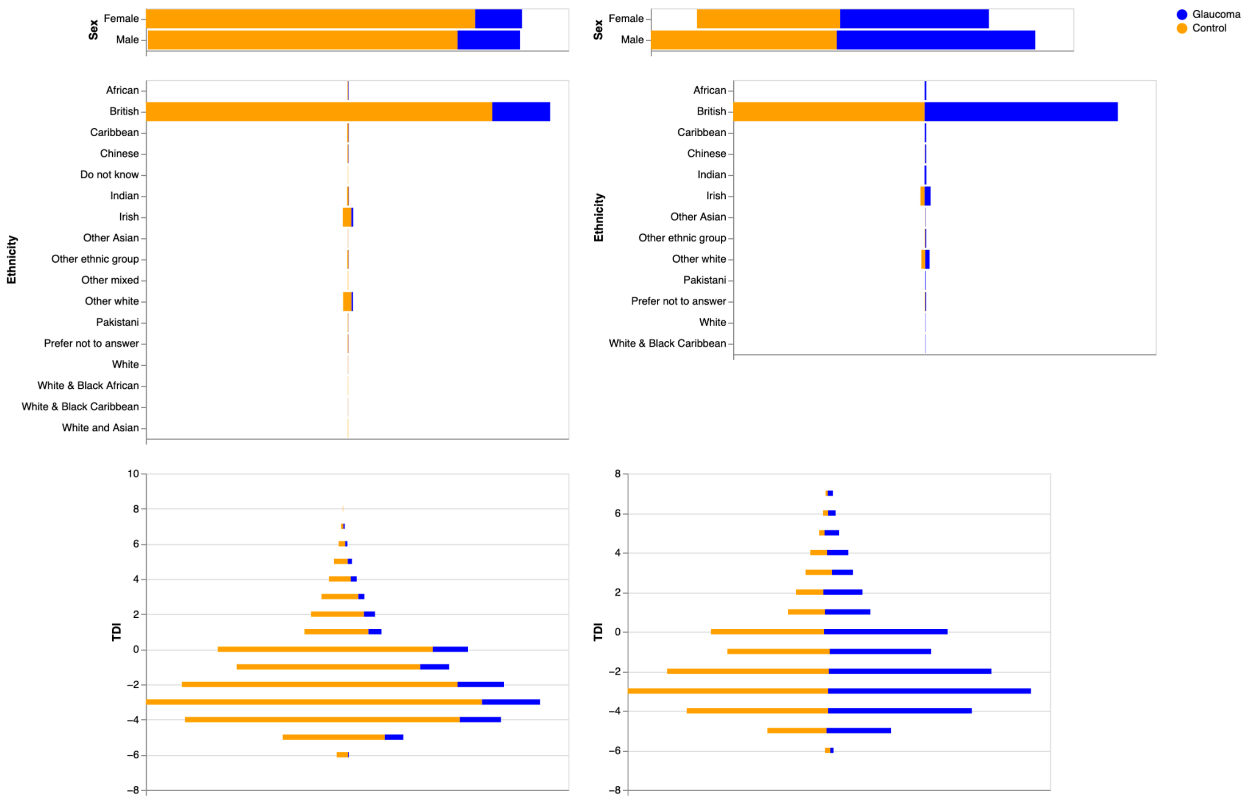


**Supplementary Figure 1.** *Distributions of sex, ethnicity, and TDI of subjects, colored by glaucoma status. The left panels show the entire dataset before matching. The right panel shows the dataset after matching each subject with glaucoma to a control subject with the same or similar age, sex, ethnicity, and TDI.*

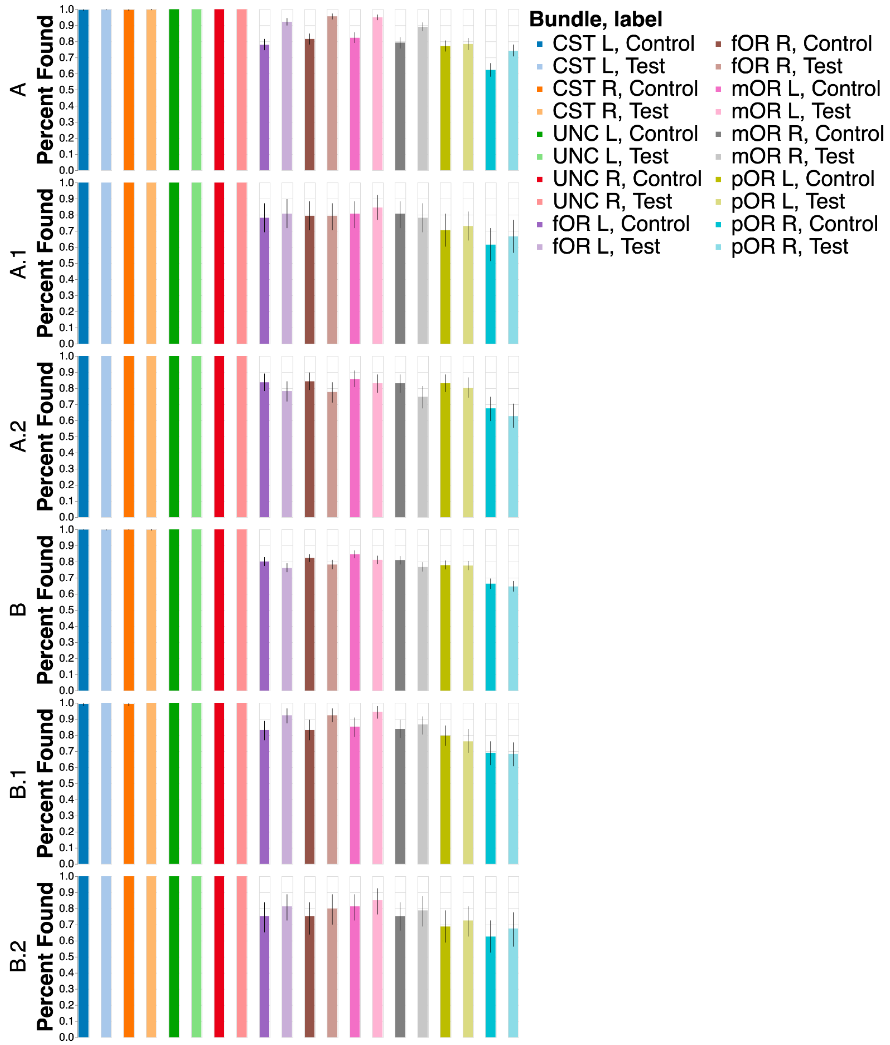


**Supplementary Figure 2.** *Percentage of successfully delineated bundles in each dataset, separated by classification label. The OR sub-bundles are harder to track than the controls, and are sometimes not found. Uncertainties show a bootstrapped 95% confidence interval. Note that, in dataset A, the control subjects are not as likely to have a successfully delineated OR as the glaucoma subjects, but see control analysis in Supplementary Figure 3.*

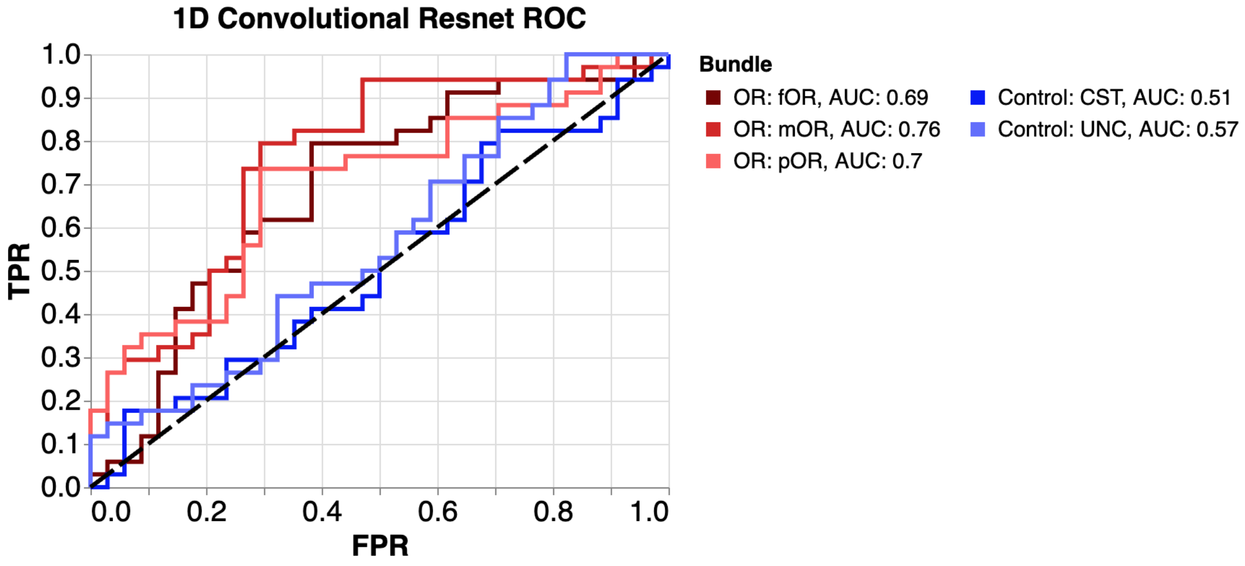


**Supplementary Figure 3.** *ROC curves for prediction of glaucoma using the five neural networks, each trained using tissue property from a different bundle. Only subjects with no missing bundles are included. Similar to in Figure 3, the OR bundles in red tend to have better performance than the control bundles. However, because fewer subjects are used, the differences in the AUC are no longer statistically significant, even though a similar effect is seen.*
